# Mutations in *RPL3L* and *MYZAP* increase risk of atrial fibrillation

**DOI:** 10.1101/223578

**Authors:** Rosa B. Thorolfsdottir, Gardar Sveinbjornsson, Patrick Sulem, Stefan Jonsson, Gisli H. Halldorsson, Pall Melsted, Erna V. Ivarsdottir, Olafur B. Davidsson, Ragnar P. Kristjansson, Gudmar Thorleifsson, Anna Helgadottir, Solveig Gretarsdottir, Gudmundur Norddahl, Sridharan Rajamani, Bjarni Torfason, Atli S. Valgardsson, Jon T. Sverrisson, Vinicius Tragante, Folkert W. Asselbergs, Dan M. Roden, Dawood Darbar, Terje R. Pedersen, Marc S. Sabatine, Maja-Lisa Løchen, Bjarni V. Halldorsson, Ingileif Jonsdottir, David O. Arnar, Unnur Thorsteinsdottir, Daniel F. Gudbjartsson, Hilma Holm, Kari Stefansson

**Affiliations:** deCODE genetics/Amgen, Inc., Reykjavik, Iceland; School of Engineering and Natural Sciences, University of Iceland, Reykjavik, Iceland; Department of Cardiothoracic Surgery, Landspitali University Hospital, Reykjavik, Iceland; Department of Medicine, Akureyri Regional Hospital, Akureyri, Iceland; Department of Cardiology, Division Heart & Lungs, University Medical Center Utrecht, University of Utrecht, Utrecht, The Netherlands; Durrer Center for Cardiovascular Research, Netherlands Heart Institute, Utrecht, the Netherlands; Institute of Cardiovascular Science, Faculty of Population Health Sciences, University College London, London, United Kingdom; Farr Institute of Health Informatics Research and Institute of Health Informatics, University College London, London, United Kingdom; Departments of Medicine, Pharmacology, and Biomedical Informatics, Vanderbilt University Medical Center, Nashville, TN, USA; Division of Cardiology, Department of Medicine, University of Illinois at Chicago, Chicago, Illinois, USA; Center For Preventive Medicine, Oslo University Hospital and Medical Faculty, University of Oslo, Oslo, Norway; TIMI Study Group, Division of Cardiovascular Medicine, Brigham and Women’s Hospital and Harvard Medical School, Boston, MA, USA; Department of Community Medicine, UiT The Arctic University of Norway, Tromsø, Norway; Reykjavik University, Reykjavik, Iceland; Faculty of Medicine, University of Iceland, Reykjavik, Iceland; Department of immunology, Landspitali University Hospital, Reykjavik, Iceland; Department of Medicine, Landspitali University Hospital, Reykjavik, Iceland

**Author notes:** Address for correspondence <u>Kari Stefansson</u>, Address: Sturlugata 8, 101 Reykjavik, Iceland. or <u>Hilma Holm</u>. Address: Sturlugata 8, 101 Reykjavik, Iceland.

**Keywords:** Atrial fibrillation, *RPL3L*, *MYZAP*, genome-wide association study, arrhythmia, UK Biobank

## Abstract

We performed a meta-analysis of genome-wide association studies on atrial fibrillation (AF) among 14,710 cases and 373,897 controls from Iceland and 14,792 cases and 393,863 controls from the UK Biobank, focusing on low frequency coding and splice mutations, with follow-up in samples from Norway and the US. We observed associations with two missense (OR=1.19 for both) and one splice-donor mutation (OR=1.52) in *RPL3L*, encoding a ribosomal protein primarily expressed in skeletal muscle and heart. Analysis of 167 RNA samples from the right atrium revealed that the splice donor mutation in *RPL3L* results in exon skipping. AF is the first disease associated with *RPL3L* and *RPL3L* is the first ribosomal gene implicated in AF. This finding is consistent with tissue specialization of ribosomal function. We also found an association with a missense variant in *MYZAP* (OR=1.37), encoding a component of the intercalated discs of cardiomyocytes, the organelle harbouring most of the mutated proteins involved in arrhythmogenic right ventricular cardiomyopathy. Both discoveries emphasize the close relationship between the mechanical and electrical function of the heart.

## Introduction

Atrial fibrillation (AF) is the most common arrhythmia of clinical significance with an estimated number of 33.5 million individuals diagnosed with AF globally in the year 2010^1^. It is associated with increased mortality and morbidity, particularly stroke and heart failure, and is responsible for substantial health care costs^1^. AF is a complex disease that is characterized by both mechanical and electrical abnormalities of the atria that may be detected prior to diagnosis of the arrhythmia itself. The role of atrial myopathy and fibrosis in the development of AF is increasingly recognized and it has been postulated that these processes may contribute to cardioembolic stroke in the absence of arrhythmia^2^. Thus, identification of the early stages of atrial myopathy may allow for therapy to prevent progression of atrial remodelling, AF and stroke^3^.

Genome-wide association studies (GWAS), assessing primarily common sequence variants, have yielded over 30 genetic loci that associate with AF^4^. Most of the associated variants are non-coding and the causative genes remain unknown but the closest genes reveal a polygenic process, implicating transcription factors, cardiac ion channels, myocardial and cytoskeletal proteins in the pathogenesis of AF. In the pre-GWAS era, linkage mapping and candidate gene sequencing linked a number of rare sequence variants to AF, mostly in single cases or familial AF, including variants in cardiac ion channel genes^4^. These mutations explain a small proportion of AF cases, and for some, the genetic evidence is not robust.

In the past few years, through GWAS based on whole-genome sequencing, we have identified three low-frequency coding variants that associate with AF^5–8^. All three variants are in structural genes, the myosin sarcomere genes *MYH6*^5^ and *MYL4*^6,7^ and the cytoskeletal gene *PLEC*^8^. These findings support the notion of an important relationship between myocardial mechanical integrity and the development of arrhythmias.

Here, we continue our search for variants associated with AF to shed further light on the pathophysiology of this common arrhythmia. We performed an AF GWAS using data from Iceland and the UK Biobank, focusing on rare and low frequency coding and splice variants, with follow-up of the most significant variants in samples from Norway and the US.

## Results

### Mutations in *RPL3L* and *MYZAP* associate with atrial fibrillation

We performed a meta-analysis on AF including 14,710 cases and 373,897 controls from Iceland and 14,792 cases and 393,863 controls from the UK Biobank,^9^ focusing on variants annotated as having moderate or high impact on protein function (including moderate: missense, in-frame indel, splice-region, and high impact: splice-acceptor, splice-donor, frameshift, stop-gained and stop lost variants)^10^. To account for the expected impact, we applied the significance thresholds of *P* < 5.1 × 10^−8^ for moderate and *P* < 2.6 × 10^−7^ for high impact variants^11^.

We identified two novel genome-wide significant associations in the gene *RPL3L* on chromosome 16, represented by the missense variant p.Ala75Val (allele frequency 3.65% in Iceland and 3.37% in UK, combined OR: 1.19, 95% CI: 1.13-1.25, *P* = 3.4×10^−12^) and the splice-donor variant c.1167+1G>A (allele frequency 0.61% in Iceland and 0.31% in UK, combined OR: 1.52, 95% CI: 1.33-1.74, *P* = 8.2×10^−10^) associating with AF (Table 1). The two variants are not correlated (D′ = 1, r^2^ = 0.00024, Supplementary Table 2) and when conditioned on each other, both associations with AF remained (Supplementary Table 3). To further assess the relationship between *RPL3L* and AF, we tested all 15 low frequency coding variants in the gene for association with AF after conditioning on p.Ala75Val and c.1167+1G>A (significance threshold = 0.05 /15 = 0.0033, Supplementary Table 4). We observed a distinct association with the missense mutation p.Gly12Arg (allele frequency 0.79% in Iceland and 0.43% in the UK, combined OR: 1.18, 95% CI: 1.05-1.34, *P* = 0.0066, Table 1). Correlation between all three mutations are shown in Supplementary Table 2 and AF association results in Iceland for each mutation after conditioning on the other two is shown in Supplementary Table 3. The *RPL3L* gene encodes a ribosomal protein (ribosomal protein like 3L) that is primarily expressed in skeletal muscle and heart unlike the ubiquitous expression of most ribosomal proteins^12^.

**Table 1.** Meta-analysis results for atrial fibrillation mutations

|  | *MYZAP* p.Gln254Pro<br>missense |  |  | *RPL3L* p.Ala75Val<br>missense |  |  | *RPL3L* c.1167+1G>A<br>splice-donor |  |  | *RPL3L* p.Gly12Arg<br>missense |  |  |
| --- | --- | --- | --- | --- | --- | --- | --- | --- | --- | --- | --- | --- |
| **Rs name**<br>**Position (hg38)** | rs147301839<br>chr15:57632516 |  |  | rs140185678<br>chr16:1953015 |  |  | rs140192228<br>chr16:1945498 |  |  | rs146294352<br>chr16:1954118 |  |  |
| **Dataset**<br>**(cases/controls)** | **Freq**<br>**%** | **OR**<br>**(95% CI)** | **P** | **Freq**<br>**%** | **OR**<br>**(95% CI)** | **P** | **Freq**<br>**%** | **OR**<br>**(95% CI)** | **P** | **Freq**<br>**%** | **OR**<br>**(95% CI)** | **P** |
| Iceland<br>(14,710/373,897) | 1.08 | 1.38 (1.20-1.60) | 9.0×10<sup>-6</sup> | 3.65 | 1.18 (1.09-1.28) | 6.4×10<sup>-5</sup> | 0.61 | 1.37 (1.14-1.65) | 8.7×10<sup>-4</sup> | 0.79 | 1.29 (1.09-1.53) | 0.0029 |
| UK Biobank<br>(14,792/393,863) | 0.36 | 1.32 (1.10-1.57) | 0.0023 | 3.37 | 1.20 (1.13-1.27) | 1.2×10<sup>-8</sup> | 0.31 | 1.71 (1.40-2.07) | 6.9×10<sup>-8</sup> | 0.43 | 1.08 (0.90-1.28) | 0.41 |
| **Meta-analysis:**<br>**Iceland and UK** |  | **1.36 (1.21-1.52)** | **7.8×10<sup>-8</sup>** |  | **1.19 (1.13-1.25)** | **3.4×10<sup>-12</sup>** |  | **1.52 (1.33-1.74)** | **8.2×10<sup>-10</sup>** |  | **1.18 (1.05-1.34)** | **0.0066** |
| Fourier<br>(1,238/11,562) | 0.21 | 1.61 (0.46-5.69) | 0.46 | 4.88 | 0.93 (0.55-1.59) | 0.79 | 0.12 | 1.59 (0.13-20.07) | 0.72 | 0.39 | 1.52 (0.56-4.12) | 0.41 |
| Vanderbilt<br>(759/759) | 0.46 | 7.03 (1.29-38.23) | 0.024 | - | - | - | - | - | - | - | - | - |
| Norway<br>(714/698) | 0.49 | 1.14 (0.39-3.32) | 0.81 | - | - | - | - | - | - | - | - | - |
| **Combined** |  | **1.37 (1.22-1.52)** | **2.9×10<sup>-8</sup>** |  | **1.19 (1.13-1.25)** | **5.0×10<sup>-12</sup>** |  | **1.52 (1.33-1.74)** | **7.7×10<sup>-10</sup>** |  | **1.19 (1.05-1.34)** | **0.0052** |

Here we show association results for atrial fibrillation for the discovery and follow-up datasets and the joint analysis of all datasets (combined). Freq = frequency of minor allele, OR = odds ratio, 95% CI = 95% confidence interval, $P = P$ value from case control association analysis. The $P$ value for heterogeneity analysis for the *MYZAP* p.Gln254Pro joint-analysis was 0.42.

Our GWAS yielded another moderate impact variant with suggestive association with AF, the missense mutation p.Gln254Pro in the gene *MYZAP* on chromosome 15 (allele frequency 1.08% in Iceland and 0.36% in UK, combined OR: 1.36, 95% CI: 1.21-1.52, *P* = 7.8×10^−8^). To assess this mutation further, we tested it for association with AF in three additional sample sets of 2,711 cases and 13,019 controls combined, from the Further Cardiovascular Outcomes Research with PCSK9 Inhibition in Subjects with Elevated Risk (FOURIER) trial, the Vanderbilt AF Registry and from Norway. Joint analysis of all datasets yielded genome-wide significant association with AF (OR: 1.37, 95% CI: 1.22-1.52, *P* = 2.9×10^−8^, Table 1). No other coding variant in *MYZAP* associates independently with AF (Supplementary Table 5). *MYZAP* encodes myozap, myocardial zonula adherens protein, which is primarily expressed in the heart in man and its homolog in the mouse has been localized to the intercalated discs^13^. Three other moderate or high impact coding variants in the genes *MYH6*, *PLEC* and *MYL4* (recessive model), previously reported by us, were genome-wide significantly associated with AF in this dataset^5–8^.

### The p.Ala75Val missense mutation in *RPL3L* associates with electrocardiogram measures in the absence of atrial fibrillation

We have previously demonstrated that the effects of reported AF variants on ECG traits measured in sinus rhythm range from none to extensive and there is no clear relationship between AF effect sizes and effects on ECG measures^8^. For example, a sequence variant associated with AF in the sodium channel gene *SCN10A* has extensive and strong effects on ECG measures but a relatively small AF effect compared to the most significant common AF variant near *PITX2* that has minimal effect on ECG measurements (Figure 1). Figure 1 shows the effects of the *RPL3L* and *MYZAP* AF variants on ECG traits compared to the effects of 31 published AF variants. For the analysis we used 289,297 sinus rhythm ECGs from 62,974 individuals not diagnosed with AF and tested all variants for association with 122 ECG variables, some of which are correlated (Supplementary Tables 6 and 7). We used the Benjamini-Hochberg false discovery rate (FDR) procedure controlling the FDR at 0.05 at each marker to account for multiple testing. The *RPL3L* missense variant p.Ala75Val associates with measures of atrial conduction during sinus rhythm, both P wave amplitude and area, and with QRS duration. None of the other variants in *RPL3L* and *MYZAP* associate with ECG traits in sinus rhythm. When testing for association with ECG traits using all ECGs irrespective of rhythm and history of AF, p.Ala75Val in *RPL3L* associates more significantly with ECG measurements and p.Gln254Pro in *MYZAP* associated with P wave morphology, R amplitude and T wave morphology (Supplementary Figure 1).

**Figure 1.**
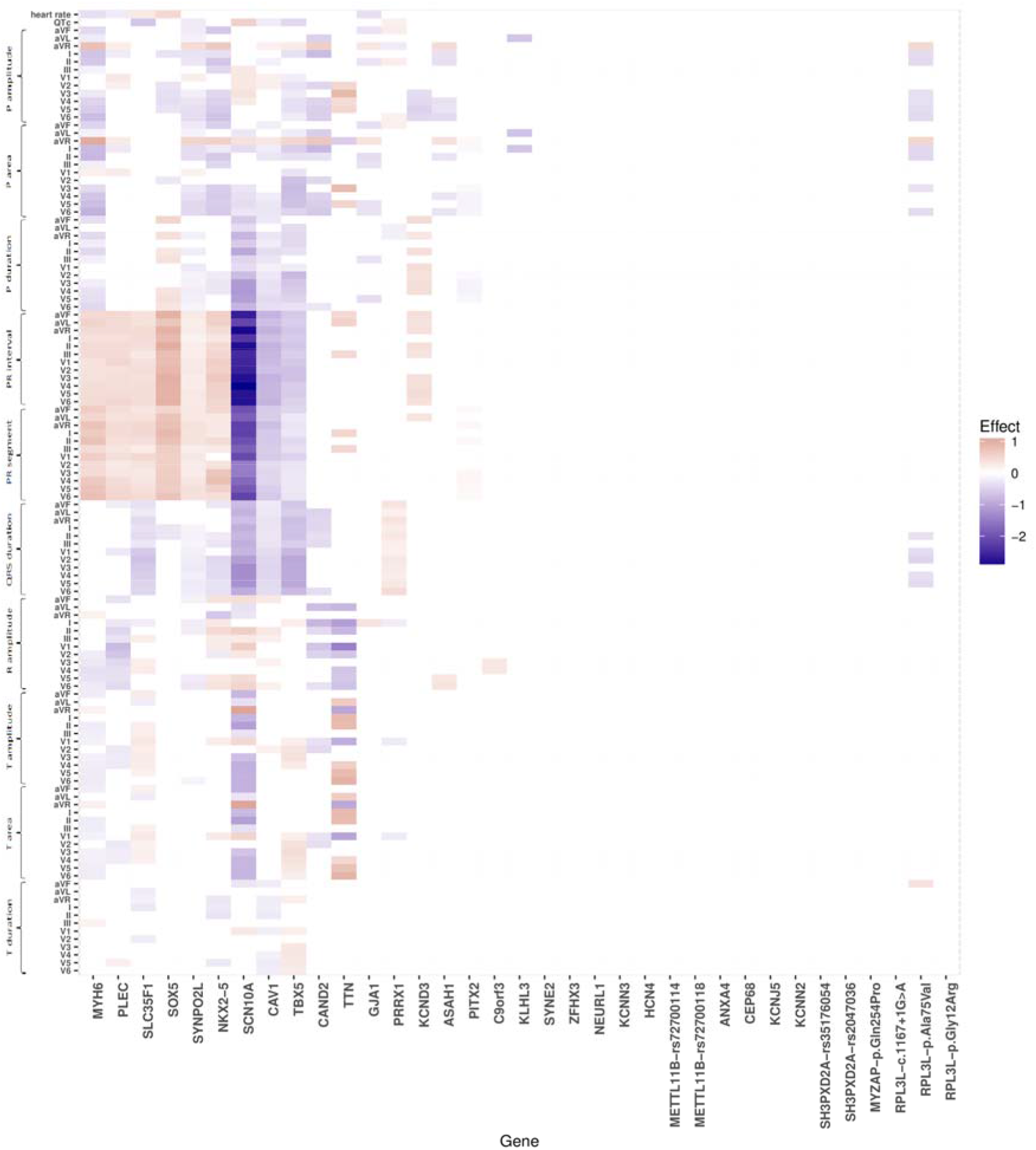
Heatmap showing the effects of atrial fibrillation (AF) variants on ECG traits in sinus rhythm ECGs, excluding AF cases. ECG measurements were available for 62,974 individuals without AF. Each column shows the estimated effect of the risk allele of an AF variant on various ECG traits. The effect of each variant, annotated with the corresponding gene name, is scaled with the log_10_-AF odds ratio. Red color represents a positive effect on the ECG variable and blue color a negative effect. The effect is shown only for significant associations after adjusting for multiple testing with a false discovery rate procedure for each variant. Non-significant associations are white in the heatmap.

### The missense mutation in *MYZAP* associates with sick sinus syndrome

Variants that associate with risk of AF also tend to associate with the related atrial arrhythmia sick sinus syndrome (SSS) and commonly with effects that are proportional to the AF effects^14^. One notable exception is the missense mutation in *MYH6* that we originally discovered through its association with high risk of SSS and confers a substantially greater risk of SSS than predicted from its effect on AF risk^5^. We tested the four new AF mutations in 3,568 SSS cases and 346,025 controls from Iceland and 403 cases and 403,181 controls from the UK Biobank.^9^ In the joint analysis, p.Gln254Pro in *MYZAP* associates with SSS (OR: 1.65, 95% CI: 1.33-2.05, *P* = 5.0×10^−6^) (Supplementary Table 8).

To gain a better understanding of the new AF mutations, we tested them for association with other phenotypes in deCODE’s genotype/phenotype database under both additive and recessive models but found no other associations passing Bonferroni correction. Association results for available relevant phenotypes including risk factors of AF are listed in Supplementary Tables 9 and 10. Since mutations in ribosomal genes are commonly associated with bone marrow failure, we specifically queried the relationship between the *RPL3L* variants and blood cells and found no associations. Similarly, since mutations in intercalated disc genes, albeit not *MYZAP*, have been associated with cardiomyopathies in man^15^ we assessed the link between the *MYZAP* mutation and cardiomyopathies in our database, but found none.

### The splice-donor mutation c.1167+1G>A in *RPL3L* causes exon skipping

We obtained RNA samples from cardiac atria of 167 Icelanders and used them to assess the effect of the splice-donor mutation c.1167+1G>A in *RPL3L*. Two of the 167 individuals carry this mutation. Non-carriers only produce the primary *RPL3L* isoform, but both carriers also produce an alternative isoform that skips exon 9 (*P* = 0.0052, Figure 2, panel a). We also found that carriers express the two isoforms in approximately equal abundance. Exon 9 is the second to last exon in *RPL3L* and is 120 base pairs long and therefore its deletion is in-frame (Figure 2, panel b).

**Figure 2.**
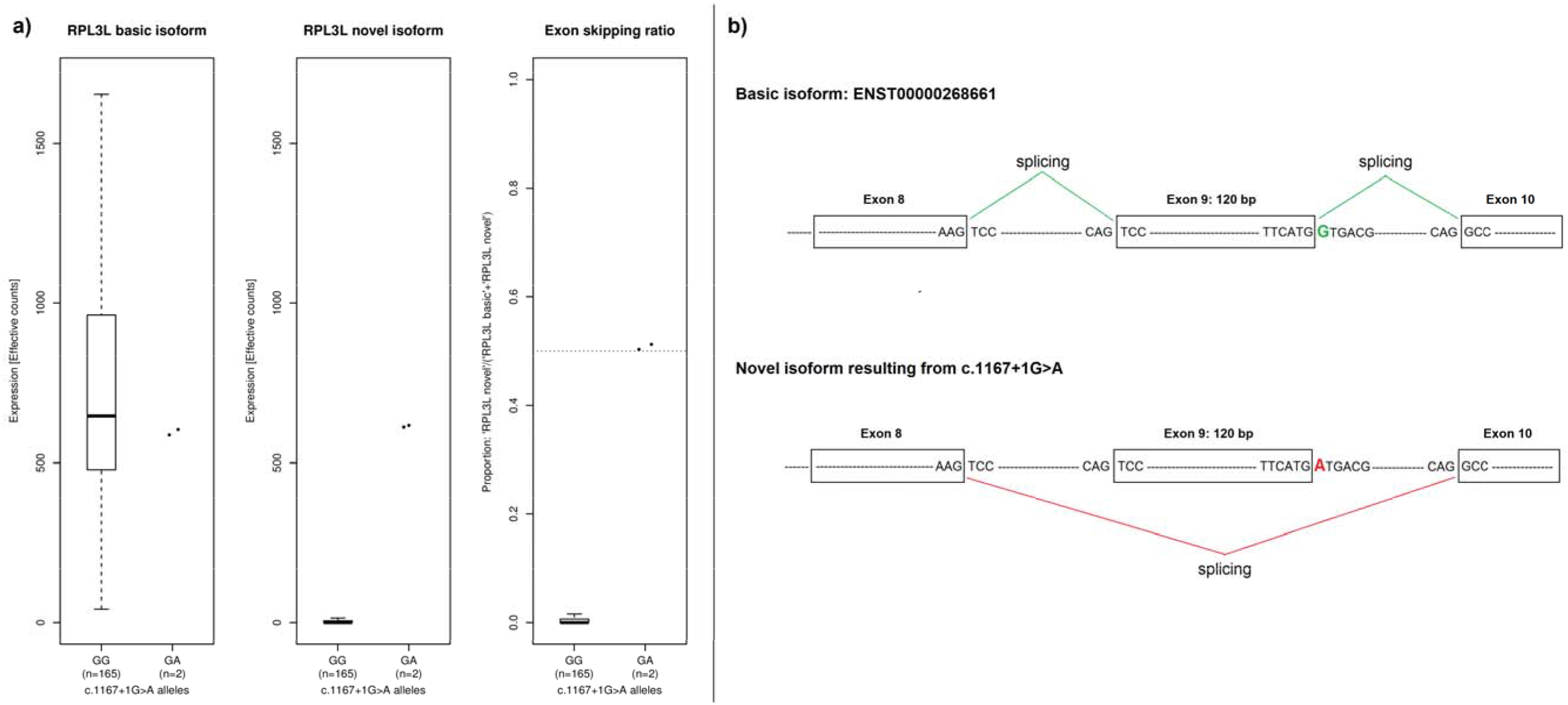
The effect of the splice-donor mutation c.1167+1G>A in *RPL3L* on splicing. *Panel a* shows quantification of two forms of RPL3L transcripts; the primary isoform, ENST00000268661, and a novel isoform with skipping of exon 9 resulting from c.1167+1G>A. It also shows the proportion of novel isoform among all transcripts. A total of 167 samples, all from the right atrium, where included in the analysis. Two of those came from carriers of c. 1167+1G>A. The figure demonstrates that only the two carriers have the novel isoform with skipping of exon 9. Their exon skipping proportion is approximately 0.5 while it is zero in non-carriers. *Panel b* is a schematic illustration of the splicing of RPL3L among carriers and noncarriers of c. 1167+G>A. The variant is in a splice donor site by the second last exon and results in exon skipping. The skipped exon is 120 base pairs and therefore its deletion is in-frame.

## Discussion

By performing a meta-analysis of AF using samples from deCODE and the UK biobank, focusing on rare and low frequency coding and splice variants, we discovered four new AF variants in two genes, three in the ribosomal gene *RPL3L* and one in *MYZAP* that encodes a component of the cardiac intercalated discs. Risk of AF has not been associated with a ribosomal gene before.

The eukaryotic ribosome, composed of four different ribosomal RNAs and ~80 ribosomal proteins, is a complex cellular machine that translates messenger RNA into protein^16^. Only a few rare inherited diseases, ribosomopathies, have been specifically linked to mutations in genes encoding ribosomal proteins or ribosome biogenesis factors. They include Diamond–Blackfan anemia and Shwachman–Diamond syndrome that are characterized by a distinct set of clinical features, including bone marrow failure and/or developmental abnormalities^17^. The ribosome has generally been considered to function in a housekeeping capacity but recent studies have revealed that ribosome activity may be regulated in a cell-specific manner, for example through changes in the protein composition of the ribosome^18,19^. One example is the *RPL3L* with expression restricted to skeletal muscle and the heart^12^. Ribosomes containing RPL3L instead of its ubiquitously expressed homolog, RPL3, have altered translational activity and RPL3L has been suggested to be a negative regulator of muscle growth^20^.

Two of the three *RPL3L* mutations that we identified are missense mutations, p.Ala75Val and p.Gly12Arg. Both Ala75 and Gly12 are highly conserved in *RPL3L* and also in *RPL3* over a range of species (PROVEAN impact prediction scores > −2.5^21^) (Supplementary Table 11). Sequencing of RNA samples from cardiac atria including from carriers of the splice donor mutation, c.1167+1G>A, demonstrated that the mutation leads to skipping of *RPL3L* exon 9, the second last exon that encodes amino acid residues 350 to 389. These residues have 75% sequence identity to the corresponding RPL3 residues. In yeast it has been shown that amino acids 373–380 in RPL3, corresponding to amino acids 382-389 in human RPL3L, form a part of the contact site of the ribosomes with the signal recognition particle that targets ribosomes to the endoplasmic reticulum membrane^22^. Based on functional similarities between RPL3 and RPL3L it is therefore possible that c.1167+1G>A disrupts engagement of RPL3L containing ribosomes with the endoplasmic reticulum and thus reducing ribosomal function. Since all three *RPL3L* mutations increase the risk of AF it could be predicted, based on the suggested effect of the splice donor mutation, that the mutations are loss of, rather than gain of, function. The association of AF with a gene expressed in the atria that is involved with regulation of muscle growth is in line with the increasingly recognized tight link between mechanical myocardial integrity and the electrical function of the heart.

The *MYZAP* gene was recently discovered by Seeger et al. in an effort to find new components of the intercalated discs^13^, a highly specialized cell-cell contact structure that enables mechanical, electrical and chemical communication between cardiomyocytes. Human Myozap mRNA is primarily expressed in the heart and in the mouse the protein was predominantly found at intercalated discs and sarcomeric Z-discs^13^. In vitro functional studies revealed a role in cardiac signal transduction as Myozap promotes serum response factor signaling in the nucleus^13^. A knockdown of the Myozap ortholog in zebrafish and cardiac overexpression of Myozap in the mouse both resulted in cardiomyopathy^13,23^, suggesting an important role of the protein in maintaining cardiac integrity and function.

According to PROVEAN^21^, Gln254 is conserved and the variant is predicted to be deleterious (Supplementary Table 11). The variant is located at the edge of the Myozap protein region associated with both activation of serum response factor-dependent transcription and actin colocalization (amino acids 91-250), and could therefore potentially affect either one or both of these protein functions^13^. An introduction of proline, a conformationally constrained amino acid, can lead to perturbations in local folding and therefore might interrupt the function of adjacent domains.

Mutations in intercalated disc genes cause cardiomyopathies, in particular arrhythmogenic right ventricular cardiomyopathy (ACM), characterized by a notable risk of both atrial fibrillation and ventricular arrhythmias, and one of the leading causes of sudden cardiac death in young people and athletes^24^. Interestingly, conduction abnormalities and arrhythmias in ACM are commonly encountered before the appearance of structural defects^15^. AF variants have also been identified in and close to genes encoding components of intercalated discs^4^, and the AF-associated gene *PITX2* has been shown to directly regulate intercalated disc genes^25^. P.Gln254Pro does not associate with cardiomyopathies, ventricular arrhythmias or sudden cardiac death in our data, suggesting that it only affects the atria but we may lack power to identify a ventricular effect.

Like p.Gln254Pro in *MYZAP*, the three low frequency missense and frameshift variants we have previously reported to increase the risk of AF, in *MYH6*, *MYL4* and *PLEC*, also increase the risk of SSS^8^. Like *MYZAP*, all three genes encode structural components of the cardiomyocyte. In particular, *PLEC* encodes a multidomain cytoskeletal linking protein which, among other functions, connects with elements of the intercalated disc and has a role in maintaining its integrity^26,27^.

In summary, we report the association of four low frequency coding variants in *RPL3L* and *MYZAP* with increased risk of AF. Using RNA samples from cardiac tissue we show that a splice-donor variant in *RPL3L* causes exon skipping. These results add to previous discoveries of three low frequency coding variants in structural genes associating with AF and highlight the intricate connection between myocardial structure and arrhythmogenesis. The association of a missense variant in *MYZAP* with AF and SSS emphasizes the role of the intercalated discs in arrhythmogenesis. The fact that a coding variant in a ribosomal protein specifically expressed in skeletal muscle and the heart increases risk of AF is in line with the novel concept of ribosome specialization in muscle and underscores the importance of this specialization for normal function of the heart. GWAS have linked a number of common variants with risk of AF but emerging discoveries of low frequency coding variants associating with AF continue to shed new light on the pathogenesis of the disease.

## Online Methods

Detailed methods are available in the Supplementary Appendix. The study was approved by the Icelandic Data Protection Authority and the National Bioethics Committee of Iceland (no. VSNb2015030021). The study complies with the Declaration of Helsinki.

### Study populations

The Icelandic AF sample consisted of 15,552 Icelanders diagnosed with AF (International Classification of Diseases (ICD) 10 code I.48 and ICD 9 code 427.3) according to electronic medical records (EMR) at Landspitali, The National University Hospital, in Reykjavik, Iceland, and Akureyri Hospital, the two largest hospitals in Iceland, between 1987 and 2017. 14,710 out of the 15,552 cases had genotypes and were included in the analysis. Controls were 373,897 Icelanders recruited through different genetic research projects at deCODE genetics. The AF population from UK Biobank consisted of 14,792 cases and 393,863 controls, all individuals of European ancestry recruited between 2006 and 2010.^9^ The UK Biobank project is a large prospective cohort study of ~500,000 individuals from across the United Kingdom, aged between 40-69 at recruitment.^9^ AF was ascertained based on ICD diagnoses. These are primary or secondary diagnoses codes a participant has had recorded across all their episodes in hospital. Diagnoses are coded according to ICD-9 or ICD-10. Self reported diagnoses were excluded from our analysis. We followed up the novel AF variants in AF sample sets from the Further Cardiovascular Outcomes Research With PCSK9 Inhibition in Subjects With Elevated Risk (FOURIER) trial^28^ (1,238 cases and 11,562 controls), the Vanderbilt AF Registry (764 cases and 762 controls) and a Norwegian AF study population from the Tromsø study (714 cases and 698 controls) (see Methods section in Supplementary Appendix). The AF variants were tested for association with other phenotypes in the deCODE genetics phenotype database which contains extensive medical information on various diseases and other traits. We also assessed the association of novel AF variants with sick sinus syndrome (SSS) among 403 cases and 403,181 controls in the UK Biobank.

### Electrocardiogram data

Electrocardiograms (ECGs) obtained in Landspitali, The National University Hospital, the largest and only tertiary care hospital in Iceland, have been digitally stored since 1998. We have analysed 434,000 ECGs obtained between 1998 and 2015. To assess the effect of AF variants on ECG traits, and thus cardiac electrical function, in the absence of AF, we excluded ECGs from individuals with AF and pacemakers and used 289,297 sinus rhythm ECGs for the primary analysis.

### Generation of genotype data

In Iceland, we identified 32.5 million sequence variants by sequencing the genomes of 15,220 Icelanders using Illumina standard TruSeq methodology to a mean depth of 35X (SD 8X) and imputed them into 151,677 chip-typed individuals and their relatives^7^ (see Methods section in Supplementary Appendix). In the UK Biobank, genotyping was performed using a custom-made Affymetrix chip, UK BiLEVE Axiom^29^, in the first 50,000 participants, and with Affymetrix UK Biobank Axiom array in the remaining participants^30^; 95% of the signals are on both chips. Imputation was performed by Wellcome Trust Centre for Human Genetics using a combination of 1000Genomes phase 3^31^, UK10K^32^ and HRC^33^ reference panels, for up to 92,693,895 SNPs^34^.

### Statistical analysis

We performed a meta-analysis on 14,710 AF cases and 373,897 controls from Iceland and 14,792 AF cases and 393,868 controls from the UK Biobank. We used logistic regression to test for association between SNPs and AF and other phenotypes in the Icelandic study, treating phenotype status as the response and allele count as a covariate. We used allele counts from genotyping or integrated over possible genotype counts based on imputation. Other available individual characteristics that correlate with phenotype status were also included in the model as nuisance variables. In Iceland these covariates were: sex, county of birth, current age or age at death (first and second order terms included), blood sample availability for the individual and an indicator function for the overlap of the lifetime of the individual with the time span of phenotype collection. In the UK biobank study 40 principal components were used to adjust for population stratification and age and sex were included as covariates in the logistic regression model. Only white British individuals were included in the study. For the meta-analysis we used a fixed-effects inverse variance method^35^ based on effect estimates and standard errors from the Icelandic and the UK Biobank study. Only sequence variants from the *Haplotype Reference Consortium* panel (HRC)^33^ were included in the meta analysis and variants from deCODE and the UK Biobank imputation were matched on position and alleles. For each study we accounted for inflation in test statistics due to cryptic relatedness and stratification by applying the method of linkage disequilibrium (LD) score regression^36^. The estimated correction factor for AF based on LD score regression was 1.39 for the additive model in the Icelandic sample and 1.04 in the UK Biobank. We corrected the threshold for genome-wide significance for multiple testing with a weighted Bonferroni adjustment using as weights the enrichment of variant classes with predicted functional impact among association signals (see Methods section of the Online Appendix for significance thresholds for specific groups of variants)^11^.

We tested AF variants for association with 122 ECG measurements using linear regression, treating the ECG measurement as the response and the genotype as the covariate. The ECG measurements were adjusted for sex, year of birth and age at measurement and were subsequently standardized to have a normal distribution. For individuals with multiple ECG measurements, the mean standardized value was used. The Benjamini-Hochberg false discovery rate (FDR) procedure controlling the FDR at 0.05 at each marker was used to account for multiple testing.

### Expression analysis in cardiac tissue

RNA sequencing was performed on samples from cardiac right atrium of 167 Icelanders (see Supplementary Table 1, for subject characteristics). The samples were obtained during cardiothoracic surgery at Landspitali, The National University Hospital, in Reykjavik, Iceland. In the case of the splice-donor mutation in *RPL3L* (c.1167+1G>A), the RNA samples from cardiac atria were used to identify a novel isoform and quantify expression at the transcript level (see Supplementary Appendix for detailed methods).

## Acknowledgements

We thank all the study subjects for their valuable participation as well as our colleagues that contributed to data collection, sample handling and genotyping. This research has been conducted using the UK Biobank Resource under Application Number ‘24711’. Dr. Darbar is supported by the NIH R01 HL092217, HL092217-07S1, R01 HL124935 and R01HL138737 grants.

## Competing financial interests

The following authors affiliated with deCODE Genetics/Amgen, Inc. are employed by the company: Rosa B. Thorolfsdottir, Gardar Sveinbjornsson, Patrick Sulem, Stefan Jonsson, Gisli H. Halldorsson, Pall Melsted, Erna V. Ivarsdottir, Olafur B. Davidsson, Ragnar P. Kristjansson, Gudmar Thorleifsson, Anna Helgadottir, Solveig Gretarsdottir, Gudmundur Norddahl, Sridharan Rajamani, Vinicius Tragante, Bjarni V. Halldorsson, Ingileif Jonsdottir, David O. Arnar, Unnur Thorsteinsdottir, Daniel F. Gudbjartsson, Hilma Holm and Kari Stefansson. Terje R. Pedersen has received consulting and/or speakers honoraria from Amgen, Sanofi, and Merck (all minor). Marc S. Sabatine has received research grant support through Brigham and Women’s Hospital from Amgen, AstraZeneca, Daiichi-Sankyo, Eisai, GlaxoSmithKline, Intarcia, Janssen Research and Development, Medlmmune, Merck, Novartis, Pfizer, Poxel and Takeda and consulting honoraria from Amgen, CVS Caremark, Esperion, Intarcia, Ionis, Janssen Research and Development, MedImmune and Merck. Bjarni Torfason, Atli S. Valgardsson, Jon T. Sverrisson, Folkert W. Asselbergs, Dan M. Roden, Dawood Darbar and Maja-Lisa Løchen have no relationship with industry to disclose.

